# Deletion of Tac1 gene impact kinase phosphorylation involved in signaling pathways associated with pain

**DOI:** 10.1101/2022.09.22.509023

**Authors:** Jennifer Ben Salem, Ji Zhang, Francis Beaudry

## Abstract

Pain in elderly persons is often not adequately treated, and current treatments may lead to poor outcomes. Therefore, new treatment strategies need to be developed based on a better understanding of the mechanisms underlying the development of chronic pain. Recent studies have shown that Tac1^-/-^ mice display a significant decrease in nociceptive pain responses to moderate or intense stimuli but present no phenotypic changes following light or nonpainful stimuli. Moreover, the deletion of the Tac1 gene led to a deficit of opioid peptides, which are essential to endogenous pain control mechanisms. Thus, we investigated whether Tac1^-/-^ mice show defective pain modulatory pathways by specifically profiling protein kinases in mice spinal cord using phosphoproteomics and bioinformatics. Protein phosphorylation is a key feature of the cellular regulatory mechanism, and phosphorylation status is related to the regulation and modulation of protein–protein binding. Bioinformatics analysis revealed that MAPK, tyrosine kinase, senescence, interleukin signaling, and TCR signaling are modulated in Tac1^-/-^ mice. Interestingly, these processes are intimately linked with inflammatory responses leading to the release of cytokines and chemokines implicated in the interactions and communications between cells. They are key players involved in the initiation and persistence of pathologic pain. The absence of the Tac1 gene products may trigger a much wider cell response to compensate for the lack of important components of the nociceptive pain transmission system.

## 1. Introduction

Pain sensation generated by noxious stimuli does not always produce the same reaction since numerous factors influence the neurophysiology of pain transmission. The nervous system encompasses various complex mechanisms that control the way noxious sensory information is perceived (Basbaum et al., 2009). It has been extensively shown that substantial modulation of sensory information occurs in the spinal cord (Honore et al., 2000; Levine et al., 1993). There are many molecular mechanisms contributing to the transmission of sensory information during the first synapse, and several key neuropeptides have been identified, including tachykinins and calcitonin gene-related peptide (CGRP) (Felippotti et al., 2012; Kuner, 2010; Mika et al., 2011). Primary sensory neurons express several peptides that act as neurotransmitters or neuromodulators. Tachykinins have been reported to play an important role in nociceptive transmission from the peripheral nervous system to the central nervous system (Khawaja and Rogers, 1996). The three main mammalian tachykinins are substance P (SP), neurokinin A (NKA) and neurokinin B (NKB), which exert their actions via interaction with a distinct set of receptors (Khawaja and Rogers, 1996; Otsuka and Yoshioka, 1993). Neuropeptides are derived from larger protein precursors recognized as proneuropeptides. The tachykinin precursor 1 gene (Tac1) encodes the protachykinin-1 protein containing the sequence of four tachykinin peptides, including SP (Cao et al., 1998). The protachykinin-1 protein is cleaved by the action of specific proteases into active neuropeptides by posttranslational proteolytic processing during axonal transport (Hook et al., 2008). SP plays an essential role in nociceptive transmission (Gao and Peet, 1999; Pailleux et al., 2013). It is a pronociceptive peptide and agonist of NK1R located in lamina I of the spinal cord (Teodoro et al., 2013; Yu et al., 1999). SP is mainly synthesized in neurons and has a widespread distribution in both the central and peripheral nervous systems. Notably, a large proportion of primary afferent neurons located in the dorsal root ganglia express a high concentration of SP, which is then transported to both the peripheral and central terminals. Additionally, the intensity, frequency, and duration of pain correlate highly with the detected concentration of SP and the expression of NK1R in the spinal cord (Sluka et al., 1997). Agonists of NK1R (e.g., SP, NKA) produce a sustained slow depolarization that contributes to the development of secondary hyperalgesia (Baumbauer et al., 2009; Dickenson, 1995; Levine et al., 1993).

Tac1^-/-^ mice displayed a significant decrease in nociceptive pain responses to moderate or intense stimuli (Cao et al., 1998). However, Tac1^-/-^ mice presented a behavioral phenotype similar to that of wild-type mice following light or nonpainful stimuli (Zimmer, 1998). Our previous study revealed that the absence of SP in Tac1^-/-^ mice hindered the release of endogenous opioid peptides, thereby negatively impacting endogenous pain-relieving mechanisms (Saidi and Beaudry, 2015). Furthermore, we have demonstrated that TAC1 is processed by proprotein convertase 1/3 (PC1/3) and proprotein convertase 2 (PC2), leading to a precursor of SP (Saidi et al., 2015). Thus, PCs could possibly be a drug target that leads to a reduction in SP biosynthesis and inhibits the development of secondary hyperalgesia. However, the partial inhibition of TAC1 processing can negatively impact the endogenous opioid system, as shown in Tac^-/-^ mice. The impairment of the endogenous opioid system would have a significant impact on patients suffering from persistent low to moderate pain and on their quality of life. Thus, we wanted to verify whether Tac1^-/-^ mice show defective pain modulatory pathways by specifically profiling protein kinases. Several kinases are implicated in peripheral and central sensitization, and their inhibition results in the attenuation of inflammatory and neuropathic pain (Khan et al., 2021; Ma and Quirion, 2005; Ruan et al., 2018).

## 2. Method

### 2.1 Sample preparation

Spinal cord tissues (n=6 per genotype) from male wild-type (C57BL/6 J) and male Tac1^−/–^ mice (product # 004103) were obtained from The Jackson Laboratory (Bar Harbor, Maine, USA) and kept frozen at –80 °C until analysis. All mice were 8 weeks old at the time of tissue collection. The animals from both groups were euthanized with an overdose of isoflurane followed by transection of the cervical spine. A flush of saline was performed within the spinal canal to extract the spinal cord, and the lumbar enlargement was collected. The tissue sample was snap-frozen in cold hexane (∼ -60 °C) and stored immediately at -80 °C pending analyses. The study protocol was approved by the Institutional Animal Care and Use Committee of the Faculty of Veterinary Medicine of the Université de Montréal, and it was performed in accordance with the guidelines of the Canadian Council on Animal Care.

Spinal cord tissues were homogenized in lysis buffer (8 M urea, 100 mM TRIS–HCl) at pH 8 at a ratio of 1:5 (w:v), and a cOmplet™ protease inhibitor tablet (Roche) was added. Lysing and homogenization were performed with a Fisher Bead Mill Homogenizer set at 5 m/s for 60 s and repeated three times with a 30-s delay. The homogenates were centrifuged at 9,000×g for 10 min. The protein concentration for each homogenate was determined using a Bradford assay(ThermoScientific, 2022). The protein concentration was adjusted to 1 mg/mL.

### 2.2 High-throughput ELISA-based antibody array method

Spinal cord homogenate samples were then analyzed using a Kinex KAM-850 antibody microarray (Kinexus Bioinformatics Corporation, Vancouver, BC, Canada). The Kinex KAM-850 antibody microarray includes 854 antibodies targeting different cell signaling pathways, including 517 pan-specific antibodies and 337 phosphosite-specific antibodies. Specifically, it included 466 antibodies against protein kinases, 44 antibodies against protein phosphatases, 46 antibodies against stress response proteins, 75 antibodies against proteins implicated in transcription and 223 others. Data are expressed as a chemiluminescence signal ratio versus the WT mouse spinal cord homogenates. The percentage change from the control (% CFC) was calculated for each protein by dividing the treated condition Z-ratio by the control Z-scores X 100 as previously described (Cheadle et al., 2003).

### 2.3 Bioinformatics

Proteins with a Z-ratio ≥ 1.2 and % error ≤ 30 were used for bioinformatics analysis. Gene Ontology analysis of genes (i.e., GO terms) was performed using Metascape (Zhou et al., 2019), where two sets of proteins (e.g., accession number) were used. The results from the upregulated and downregulated protein analyses were used to create bar plots in PRISM using Metascape reports with the - log_10_ p value of GO process matches given by Metascape. Further pathway analysis was performed using Metascape, ClueGO (Bindea et al., 2009) and Cytoscape (Shannon et al., 2003) using the REACTOME pathway databases. Significant pathways for proteins with a Z-ratio ≥ 1.2 and % error ≤ 30 were identified, comparing the ratio of target genes identified in each pathway to the total number of genes within the pathway. The statistical test used to determine the enrichment score for KEGG or REACTOME pathways was based on a right-sided hypergeometric distribution with multiple testing correction (Benjamini and Hochberg, 1995).

## 3. Results and discussion

Protein phosphorylation is a key feature of the cellular regulatory mechanism since several proteins, enzymes and receptors are either activated or deactivated by phosphorylation and dephosphorylation reactions catalyzed by kinases and phosphatases (Ardito et al., 2017). Furthermore, most signaling pathways implicate a large set of protein–protein interactions, and phosphorylation status is strongly associated with the regulation and modulation of protein– protein binding. As a result of mutations (e.g., disease-related, radiation, chemicals, and pathogens) or genetic manipulations, losses of phosphorylation sites might disrupt protein binding and deregulate signal transduction, including pain signaling pathways. As we have shown previously, the absence of SP, a proteolytic product of TAC1, has an impact on the abundance of opioid peptides and therefore on the endogenous pain-relieving mechanisms (Saidi and Beaudry, 2015). Thus, we wanted to verify whether Tac1^-/-^ mice show defective pain modulatory pathways by specifically profiling specific protein kinases.

We performed phosphoproteomics using the Kinex KAM-850 antibody microarray. Figure 1A displays up- and downregulated site-specific phosphoproteins (i.e., Z-ratio ≥ 1.2 and % error ≤ 30) using a heatmap feature. A total of 29 phospho-specific sites were either phosphorylated (e.g., upregulated) or dephosphorylated (e.g., downregulated) in Tac^-/-^ mice compared to WT mice. Specifically, 13 were upregulated, and 16 were downregulated. This is an interesting observation because it outlines the impact of deleting the Tac1 gene not only impaired the biosynthesis of specific neuropeptides (i.e., SP and NKA) but also potentially affect protein– protein interactions regulated or modulated by phosphorylation. Kinases are key molecules responsible for the transduction of signals involved in a number of cellular pathways and functions in response to a variety of stimuli. Abnormal functions of kinases have been recognized in several diseases including cancer, pain, inflammatory disease and diabetes(Ji et al., 2009; Lawrence et al., 2008). To better understand the impact of these observations, we performed REACTOME enrichment analyses.

**Figure 1.**
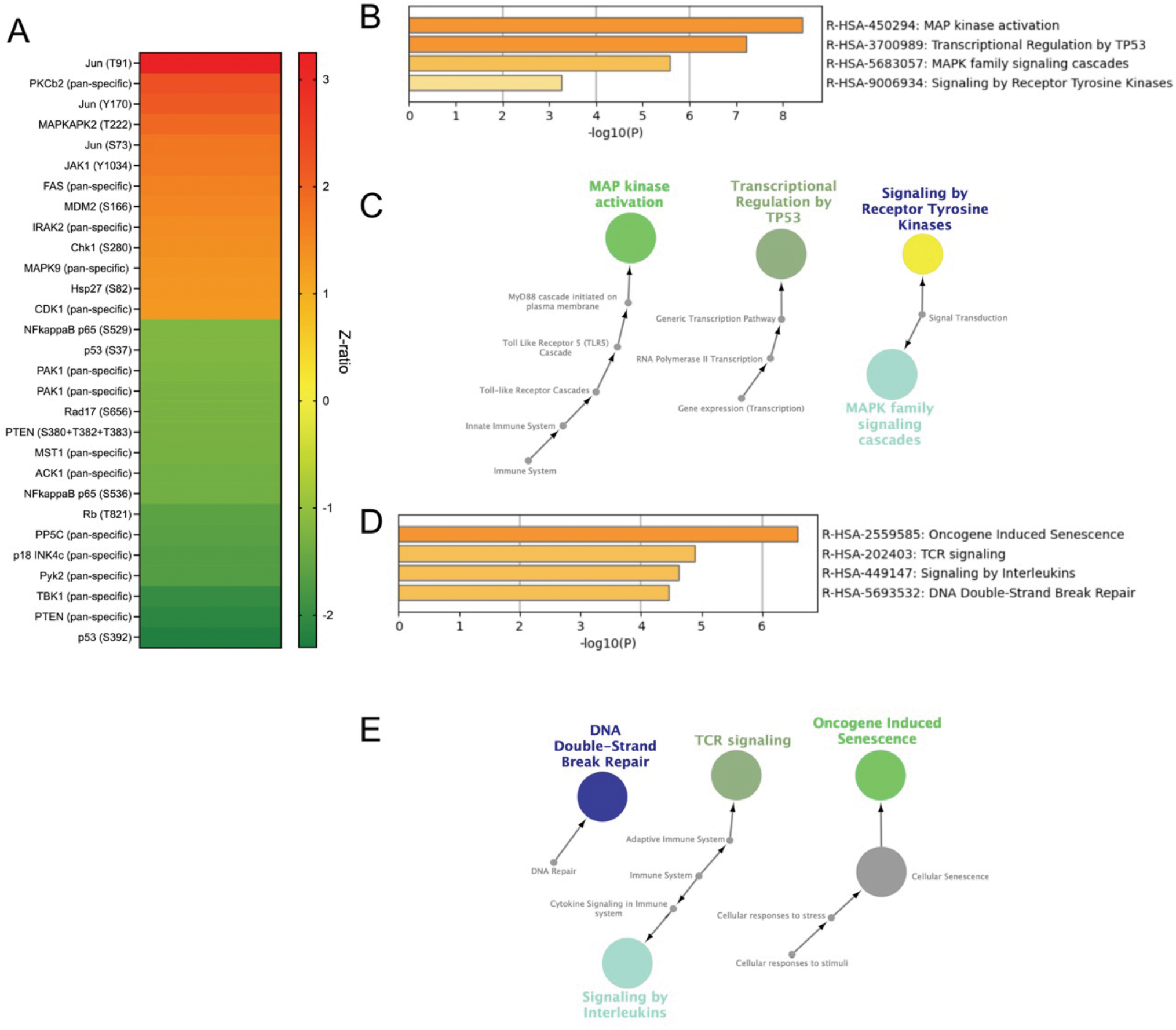
Phosphoproteomics and bioinformatics analyses for the set of proteins found to be up- or downregulated in Tac1^-/-^ spinal cord tissues. **A**. Heatmap including up- or downregulated site-specific phosphorylation. Cells are colored based on Z-ratio values. **B**. REACTOME pathway enrichment analysis for upregulated site-specific phosphorylation. **C**. Functional analysis of enriched REACTOME terms from parent node to root node for upregulated site-specific phosphorylation. **D**. REACTOME pathway enrichment analysis for downregulated site-specific phosphorylation. **E**. Functional analysis of enriched REACTOME terms from parent node to root node for downregulated site-specific phosphorylation.

As revealed in Figure 1B and C, upregulated site-specific phosphoproteins are involved in basic signaling pathways associated with pain, including mitogen-activated protein kinase (MAPK) and tyrosine kinase. MAPKs are involved in intracellular signal transduction and play fundamental roles in the regulation of neural plasticity and inflammation (Ji et al., 2009). Interestingly, a recent study revealed a reduction in nociceptive thresholds leading to enhanced hyperalgesia through the activation of MAPK-CREB signaling (Freitas et al., 2019). Tac1 KO mice have reduced nociceptive pain responses to moderate to intense stimuli but a normal response to mild stimuli. It is now well accepted that cellular tyrosine kinases play a role in the regulation of ion channel function, including TRP channels (Jin et al., 2004). These channels participate in the transduction of noxious stimuli, notably TRPV1 and TRPA1. The TP53 transcription factor has an important role in cellular responses to stress and for regulating senescence, a stable cell cycle arrest that can be triggered in normal cells in response to various intrinsic and extrinsic stimuli (Mijit et al., 2020). Senescent cells can accumulate in several tissues and contribute to the development of chronic pathologies, including chronic pain. It is strongly linked to the secretion of proinflammatory factors since senescent cells promote the production of cytokines and chemokines. Figure 1D and E show that downregulated site-specific phosphoproteins are involved in senescence, interleukin signaling, TCR signaling and DNA repair. Senescence and interleukin signaling are interrelated and potentially a response to counteract TP53 action. Chronic or neuropathic pain is frequently associated with the activation of the immune system, as depicted by a significant increase in circulating proinflammatory cytokine concentrations. Cytokine signaling between immune, glial, and neuronal cells is fundamental for the development of chronic pain. T cells, through T-cell receptors (TCRs), are involved in the regulation of the adaptive immune system and can have a pro- or antinociceptive function (Galvin and C, 2021). Therefore, the modulation of TCR signaling can impact the pain phenotype observed in Tac1 KO mice. Interestingly, MAPK signaling appears to be central to the DNA damage response (Hurley and Bunz, 2007; Ulm et al., 2001). This is an integral part of the adaptive response to stress. The downregulation of components associated with double-strand DNA break repair might be a direct response following the upregulation of components of the MAP and tyrosine kinases. All these changes are normally related to physiologic or pathologic causes. The absence of the Tac1 gene products may trigger a much wider cell response to compensate for the lack of important components of the nociceptive pain transmission system. This may have a profound impact on the phenotype observed. Future phenotypic and proteomic investigations using Tac1 KO mice and a neuropathic pain model might help to better understand why tachykinin receptor 1 (NK1) antagonists failed when tested as potential pain treatments (Hill, 2000).

## Supporting information

Kinex Antibody Microarray Data

## Acknowledgments

This project was funded by the National Sciences and Engineering Research Council of Canada (F. Beaudry discovery grant no. RGPIN-2020-05228). F. Beaudry is the holder of the Canada Research Chair in metrology of bioactive molecule and target discovery (grant no. CRC-2021-00160). This research was undertaken, partly, thanks to funding from the Canada Research Chairs Program. Ph.D. scholarships were awarded to J. Ben Salem from the Fonds de Recherche du Québec - Santé (scholarship no. 302490) and from the Université de Montréal.

## Data availability

The data that support the findings from this study are available from the corresponding author upon reasonable request.

